# The Effect of Plasma Activated Water on the Rhizosphere Composition of Arabidopsis and *Solanum lycopersicum*

**DOI:** 10.1101/2025.10.15.682576

**Authors:** Jon Kizer, Xavious Allen, Conner Robinson, Amy Grunden, Katharina Stapelmann, Marcela Rojas-Pierce

**Affiliations:** Department of Plant and Microbial Biology, North Carolina State University, Raleigh, North Carolina, United States of America; Department of Nuclear Engineering, North Carolina State University, Raleigh, North Carolina, United States of America

## Abstract

The emerging field of plasma agriculture investigates the potential benefit of non-thermal plasma (NTP) for agricultural practices. NTP-treated water, referred to as plasma activated water (PAW), has been proposed as a sustainable alternative to conventional nitrogen (N) fertilizers. Growing demand for N fertilizer is concomitant with increased global food demands. PAW contains nitrate (NO_3_^-^) and reactive oxygen species, such as hydrogen peroxide (H_2_O_2_), which are fixed from atmospheric molecules via NTP. While early studies report positive effects of PAW on plant growth, its influence on plant-associated microbial communities remains poorly understood. Here, we compared the impacts of PAW or NO_3_^-^ solutions on the rhizosphere microbial community of *Arabidopsis thaliana* and *Solanum lycopersicum*. PAW was generated by a radio frequency (RF) glow discharge plasma source and contained no measurable ROS, while the control solution contained an equivalent concentration of NO_3_^-^. No significant differences in alpha diversity were detected in either plant species microbiome after 5 weeks of treatment when grown in non-commercial potting substrate. Significant dissimilarity was found in terms of beta diversity, but the relative abundance of the sequenced genera suggested no functional differences in rhizosphere communities. Overall, PAW treatment did not adversely impact the rhizosphere microbiome in either Arabidopsis or tomato. These results support the use of PAW as an alternative N-fertilizer, though outcomes may differ for PAW solutions containing ROS.

## INTRODUCTION

Global food demand continues to rise, driving increased use of Nitrogen (N) fertilizers (Lassaletta et al., 2014). Synthetic fertilizers represent the largest portion of N-fertilizers utilized globally in agriculture (Lassaletta et al., 2014; Menegat et al., 2022). The energy-intensive Haber-Bosch process, used to fix atmospheric N, contributes 2.9 tons of atmospheric CO_2_ globally (Menegat et al., 2022; M. Wang et al., 2021), and is the primary method for industrial synthetic N fertilizer production. This reliance on unsustainable fertilizer production has spurred development of greener alternatives to generate N fertilizers while also meeting global demands (Chen et al., 2018). Plasma agriculture applies non-thermal plasmas (NTP) to fix atmospheric N_2_ into NO_3_^-^, a plant-available form. This provides a potential low-carbon alternative for fertilizer production. Treating water with NPT produces Plasma-Activated Water (PAW) or Plasma-Treated Water (PTW), a liquid fertilizer containing nitrate (NO_3_^-^). Depending on the production method, the resulting PAW solution may contain concentrations of reactive oxygen species (ROS) such as hydrogen peroxide (H_2_O_2_) and superoxide (O_2_^-^) or reactive nitrogen species (RNS) such as nitric oxide (NO) or peroxynitrite (ONOO^-^).

A healthy plant-associated rhizosphere microbiome is vital for robust plant growth as it is important for nutrient cycling and availability for plant use (Berendsen et al., 2012; Berg et al., 2014; Hein et al., 2008; Jafariyan & Zarea, 2016; Ney et al., 2019; Richardson & Simpson, 2011; Schneijderberg et al., 2020; Zhang et al., 2022). At the same time, the plant and soil environment strongly influence the diversity and function of the rhizosphere microbiome (Ai et al., 2015; Berendsen et al., 2012; Berg et al., 2014). Given that fertilizer application has major impacts on the plant and soil environment, it also affects the rhizosphere microbiome composition and performance in crop plants (Igiehon & Babalola, 2018; Shi et al., 2020; Zhu et al., 2016). For example, Zhu *et* al. reported a significant correlation between N application rate and microbial abundance in the rhizosphere. Increased N fertilizer application rates are also correlated with increased rates of exudates released from the plant host roots (Zhu et al., 2016). Additionally, since ROS play a key role in how plants respond to different environmental cues (Tsukagoshi et al., 2010), introduction of ROS and RNS from PAW (together referred to as RONS) can alter stress signaling pathways and growth or the interactions with the microbiome (Berrios & Rentsch, 2022; Jafariyan & Zarea, 2016). It is therefore important to evaluate the impact of PAW on plant-associated microbiomes. To that end, in this study, we investigate the effect of PAW without measurable ROS on rhizosphere microbiomes of Arabidopsis and tomato.

## RESULTS

### Rhizosphere Microbiome Analysis of *Arabidopsis thaliana*

Understanding whether PAW treatment alters the plant-associated microbiome is important to ascertain its potential as a N fertilizer substitute. We first tested whether PAW treatment had any effect on the Arabidopsis rhizosphere microbiome. A Radio Frequency (RF) glow discharge plasma source (Byrns et al., 2012; Lindsay et al., 2014) was used to generate PAW with 4.8 mM (310 ppm) of NO_3_^-^, which is sufficient for Arabidopsis growth (Boer et al., 2020). This PAW was specifically produced with no detectable H_2_O_2_ levels, given the negative effect of ROS-containing PAW on Arabidopsis root growth (Kizer et al., 2025).

Wild-type Arabidopsis seedlings were first germinated in sterile gel media before transplanting into substrate. The substrate was a mixture of commercial peatmoss with vermiculite and perlite not previously exposed to Arabidopsis plants, and therefore, the recruited microbes originate from the substrate itself or the seedlings (Truyens et al., 2013; Truyens et al., 2015). Plants were treated weekly with either PAW or NO_3_^-^ control solution as the only source of N. Plant roots were harvested after 5 weeks for rhizosphere microbiome analysis. DNA was extracted from rhizosphere samples and underwent 16S rRNA amplicon sequencing. The highly-variable V4 region was amplified using 515f (Parada) and 806r (Apprill) primers (Apprill et al., 2015; Parada et al., 2016), as this allows for effective classification accuracy and adequate diversity coverage with limited bias (Claesson et al., 2010; Walters et al., 2015). Amplicon sequencing of the V4 region with these primers has become a standard for sequencing of plant-associated bacteria (Gilbert et al., 2014; Walters et al., 2015), and provides adequate coverage (Breidenbach et al., 2016). Taxonomic identification was performed at the genus level to provide meaningful insight into the microbiome community structure under the conditions tested.

Both Shannon and Simpson indices were used to determine the microbial biodiversity. Alpha diversity analysis showed that both PAW-treated and NO_3_^-^-treated plants exhibited similar species richness in the rhizosphere microbial population. A Kruskal-Wallis test yielded statistically insignificant p-values at a 0.05 alpha of 0.4497 and 0.2899 for Shannon and Simpson statistics, respectively (Fig. 1). This indicated that PAW treatment did not affect overall species richness in Arabidopsis-associated rhizosphere microbial community when compared to the NO_3_^-^ control.

**Figure 1.**
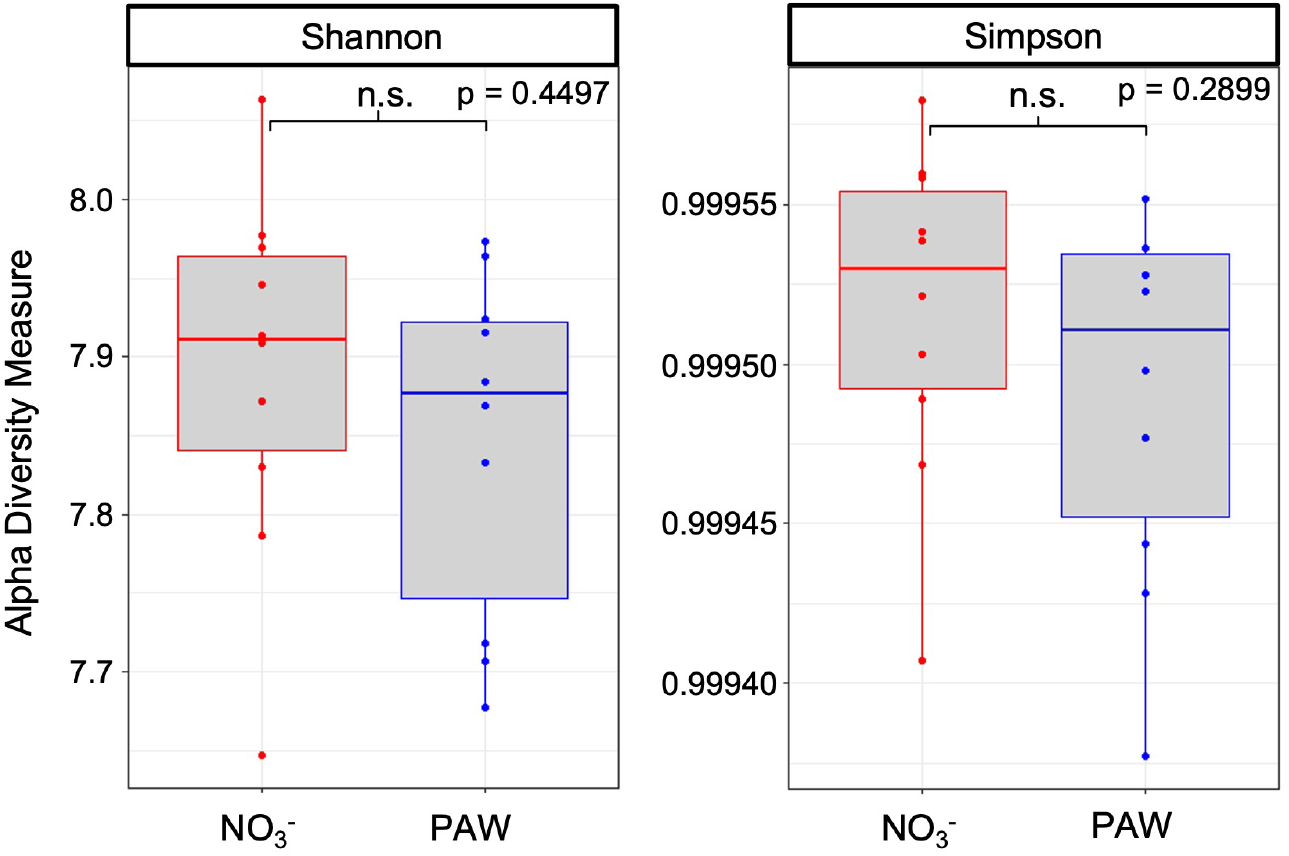
Rhizosphere microbiomes from Arabidopsis treated with PAW or the nitrate control exhibit similar taxa richness. Arabidopsis rhizosphere microbiomes were identified from plants treated with PAW containing 4.8 mM NO_3_^-^ or a control with equivalent NO_3_^-^ concentration for 5 weeks. Amplicon Sequence Variants (ASV) were derived from sequencing data and Shannon and Simpson statistics were calculated and plotted from the ASVs. A student’s T-test was performed to evaluate statistical significance. N= 10 samples per treatment.

Beta diversity of the PAW- and nitrate-treated plants was calculated using the Bray-Curtis dissimilarity method. The resulting distance statistic matrix was suitable for exploratory investigations such as the one conducted for this study (Hein et al., 2008; Schneijderberg et al., 2020). From the distance matrix, the data was plotted using Non-metric Multi-Dimensional Scaling (NMDS) and Principal Coordinate Analysis (PCoA) to scale and for visualization. The NMDS plot yielded a partial crossover indicating that there are shared microbial community members across between the two treatments (Fig. 2 A). However, further analysis and visualization with the PCoA plot showed separation of the two treatments with 9.4% and 8.2% for Axis 1 and 2, respectively (Fig. 2B). An R-package Residual Randomization in a Permutation Procedure (RRPP) analysis was then performed as a post-hoc test utilizing the distance matrix, which determined a statistically significant result at a 0.05 alpha, which indicates that there is significant dissimilarity in the rhizosphere bacteria composition between treatments. However, this variation may not be reflected in functional differences in rhizosphere microbiome performance because of functional redundancy that exists within the two Arabidopsis rhizosphere community structures.

**Figure 2.**
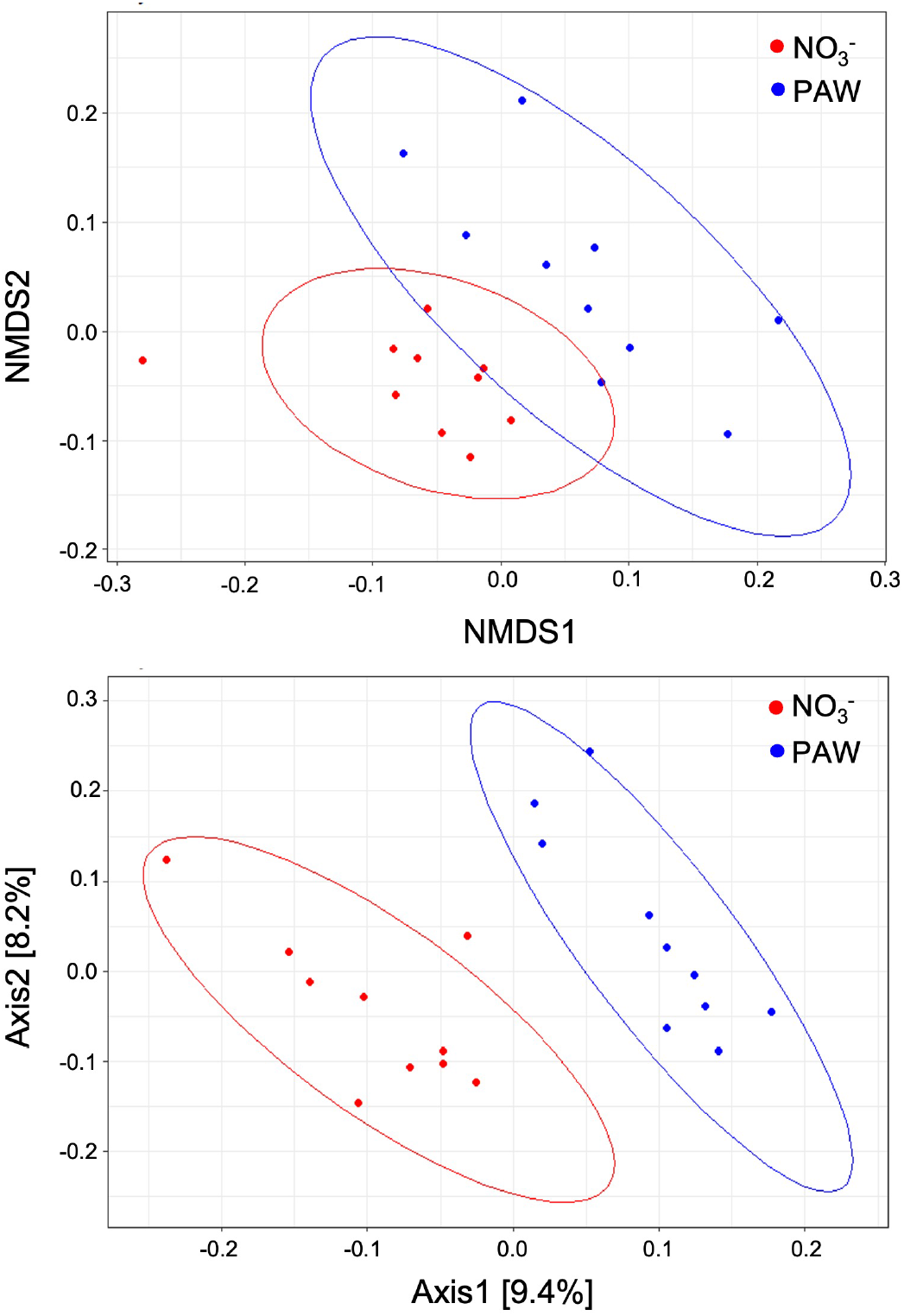
Rhizosphere microbiomes from Arabidopsis treated with PAW or the nitrate control exhibit dissimilarity in community makeup. Arabidopsis rhizosphere microbiomes were identified from plants treated with PAW (4.8 mM NO_3_^-^) or a solution of equivalent NO_3_^-^ concentration for 5 weeks. Amplicon Sequence Variants (ASV) were derived from the sequencing data. Bray-Curtis distances were calculated and plotted via Non-metric Multi-Dimensional Scaling (NMDS) (top) and a Principal Coordinate Analysis (PCoA) (bottom). A residual randomization in permutation procedures (RRPP) analysis was conducted in R to find statistical significance. N= 10 samples per treatment

Lastly, taxonomic data from the Silva bacterial 16S rRNA database was utilized to determine the identity and relative abundance of each Amplicon Sequence Variants (ASVs) in our samples at the genus taxonomic level (Quast et al., 2012). The Phyloseq R package (McMurdie & Holmes, 2013) was used for analysis, and each identified taxa was grouped by family. The top 50 taxa were then plotted and represented for each sample (Fig. 3). Interestingly, bacteria of the combined genus *Burkholderia-Caballeronia-Paraburkholderia* (BCP) were prominently represented across both treatment types (Fig. 3). This family has been associated with plant growth-promoting function (O’Sullivan & Mahenthiralingam, 2005; Suárez-Moreno et al., 2012; K. Wang et al., 2021).

**Figure 3.**
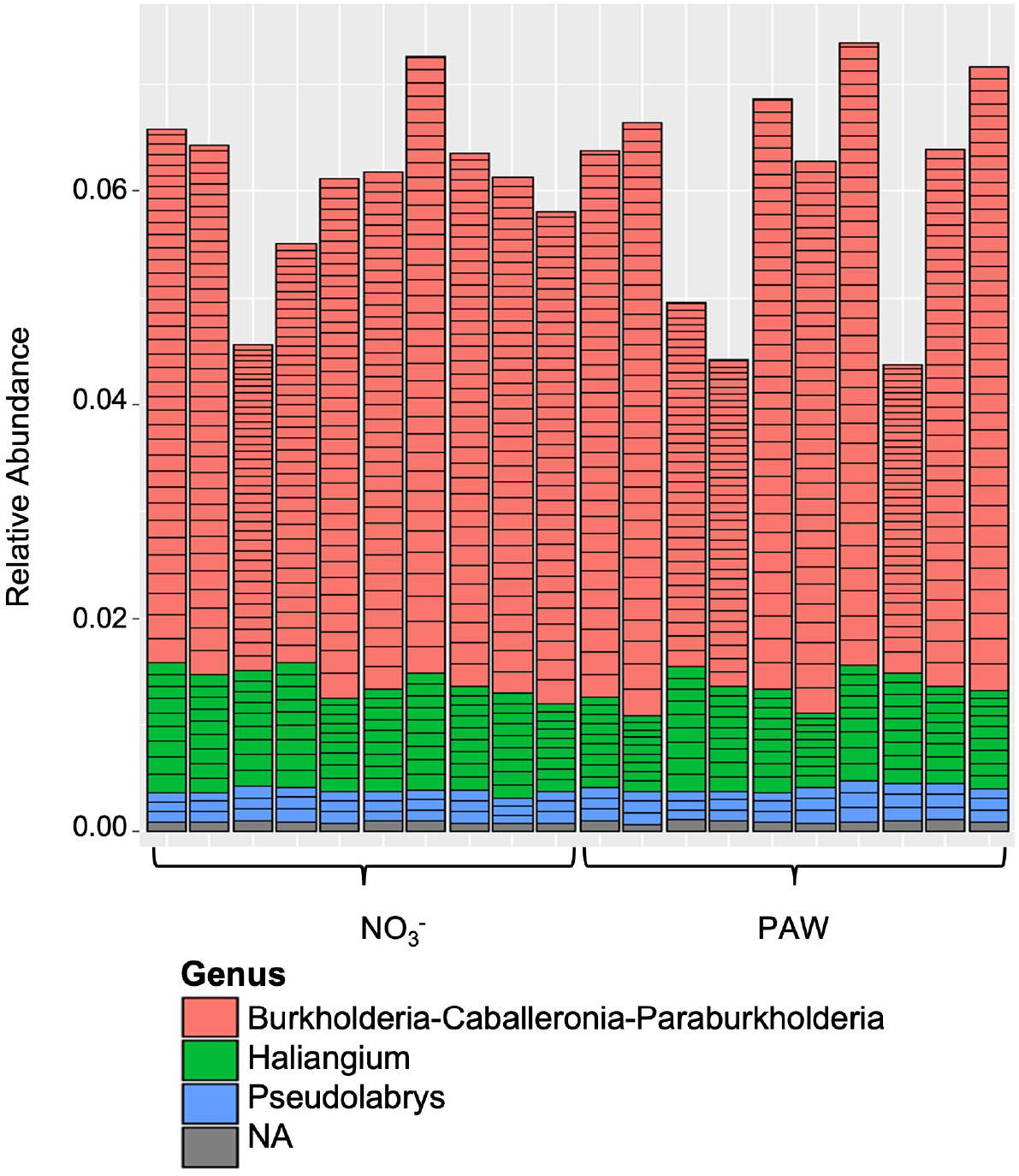
The rhizosphere microbiome from Arabidopsis treated with PAW or the control treatment exhibit similar bacterial families represented in the 50 most abundant taxa. Arabidopsis rhizosphere microbiomes were identified from plants treated with PAW (4.8 mM NO_3_^-^) or a solution of equivalent NO_3_^-^ concentration for 5 weeks and Amplicon Sequence Variants (ASV) were derived from the sequencing data. Taxonomic identities were assigned to ASVs with the Phyloseq R package. The 50 ASVs with the greatest relative abundance from each sample were plotted and grouped by genus.

### Rhizosphere Microbiome Analysis of *Solanum lycopersicum*

The effect of PAW treatment on the rhizosphere microbiome of a crop species, *S. lycopersicum*, was also investigated. PAW containing 6.5 mM (400 ppm) NO_3_^-^ and no detectable H_2_O_2_ was used given the higher N need of this crop species.

The same substrate mixture was used to control nutrients for this experiment, and seeds were directly planted into substrate. A 0.75x concentrated Hoagland solution without N was added biweekly to provide all necessary nutrients except N. Tomato seedlings were treated weekly with PAW or the NO_3_^-^ control solution, and all plants were harvested after 5 weeks. Plant height, total dry biomass, and root:shoot dry biomass ratio were measured, and no differences were detected between PAW-treated tomatoes and the NO_3_^-^ control (Supplemental Fig. 1).

Ten tomato seedlings per treatment were removed, and the substrate attached to the roots was harvested. Microbial genetic material from the rhizosphere compartment of the plant-associated microbiome was analyzed in the same manner as was performed for the Arabidopsis microbiome analysis. When comparing the rhizosphere microbiome of PAW-treated samples to that of the NO_3_^-^ control, the Shannon diversity analysis indicated significant differences (p = 0.0413), while the Simpson diversity analysis did not (p = 0.3643). Significance was calculated at a 0.05 alpha using a student’s T-test (Fig. 4). These small differences suggested limited differences in species richness between the two treatments. The Shannon index correlates to species richness and evenness, but all genera are weighed equally, causing rarer genera, which are common in microbial studies, to potentially bias the statistic (Hill et al., 2003; Hong et al., 2006; Shannon, 1948). With the Shannon index showing significance, but the Simpson showing the contrary, this implies there may be a number of taxa found only in a small number of our samples that bias the Shannon index.

**Figure 4.**
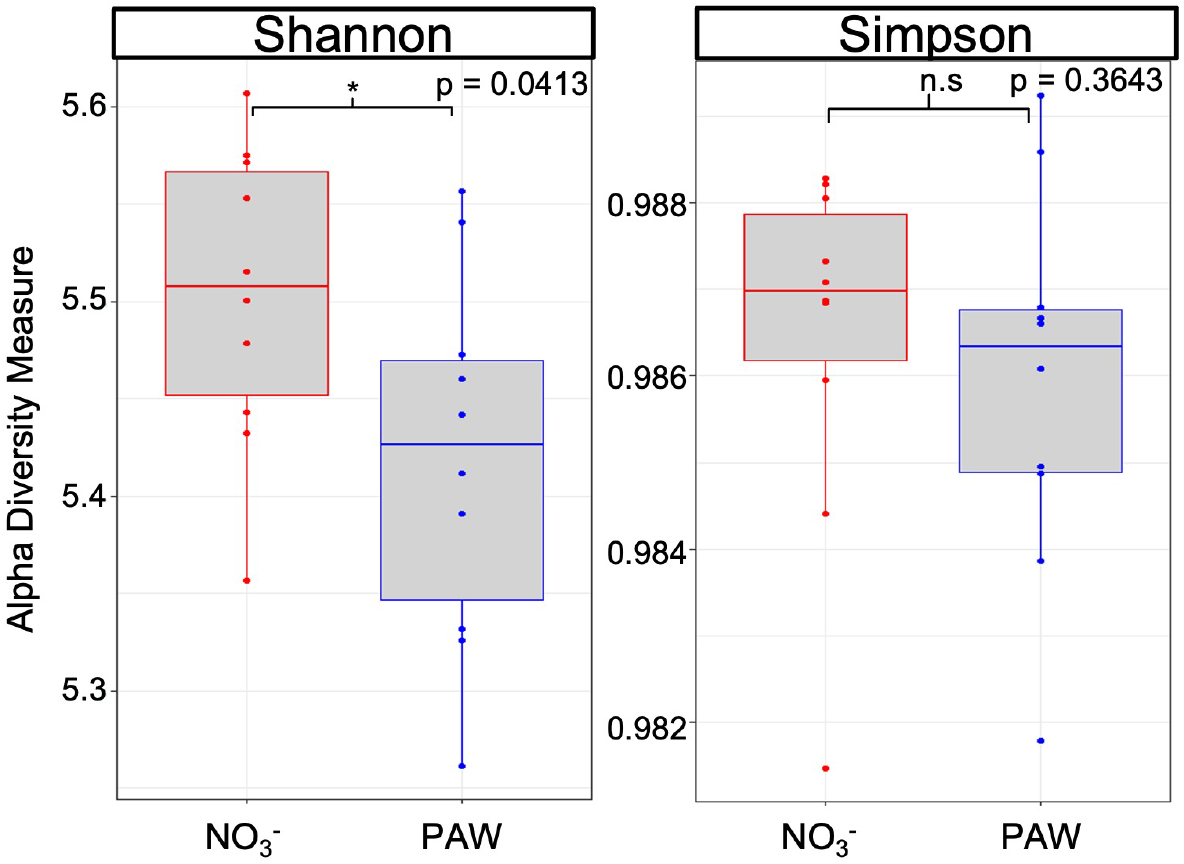
The rhizosphere microbiomes of tomato treated with PAW or the control treatment exhibit mild variation in taxa richness. Tomato plant rhizosphere microbiomes were identified from plants treated with PAW (6.5 mM NO_3_^-^) or a solution of equivalent NO_3_^-^ concentration for 5 weeks and Amplicon Sequence Variants (ASV) were derived from the sequencing data and Shannon and Simpson statistics were calculated and plotted from the ASVs. A student’s T-test was performed to evaluate statistical significance. N= 10 samples per treatment

Next, beta diversity was determined by Bray-Curtis dissimilarities between treatment groups, and the distance matrix was plotted utilizing NMDS and PCoA. The NMDS plot exhibited visual separation between the two treatments (Fig. 5A), suggesting dissimilarity in bacterial genera present between plants treated with PAW versus the NO_3_^-^ control. The PCoA plot provided visual confirmation with calculated separation with Axis1 and 2 values of 39% and 17.9%, respectively (Fig. 5B). While both treatments clustered separately, the two clusters were in close proximity. An RRPP analysis of the distance matrix was performed to confirm statistical significance. Similar to Arabidopsis, this analysis identified differences in bacterial genera that are present in the rhizosphere of PAW-treated tomato compared to the control.

**Figure 5.**
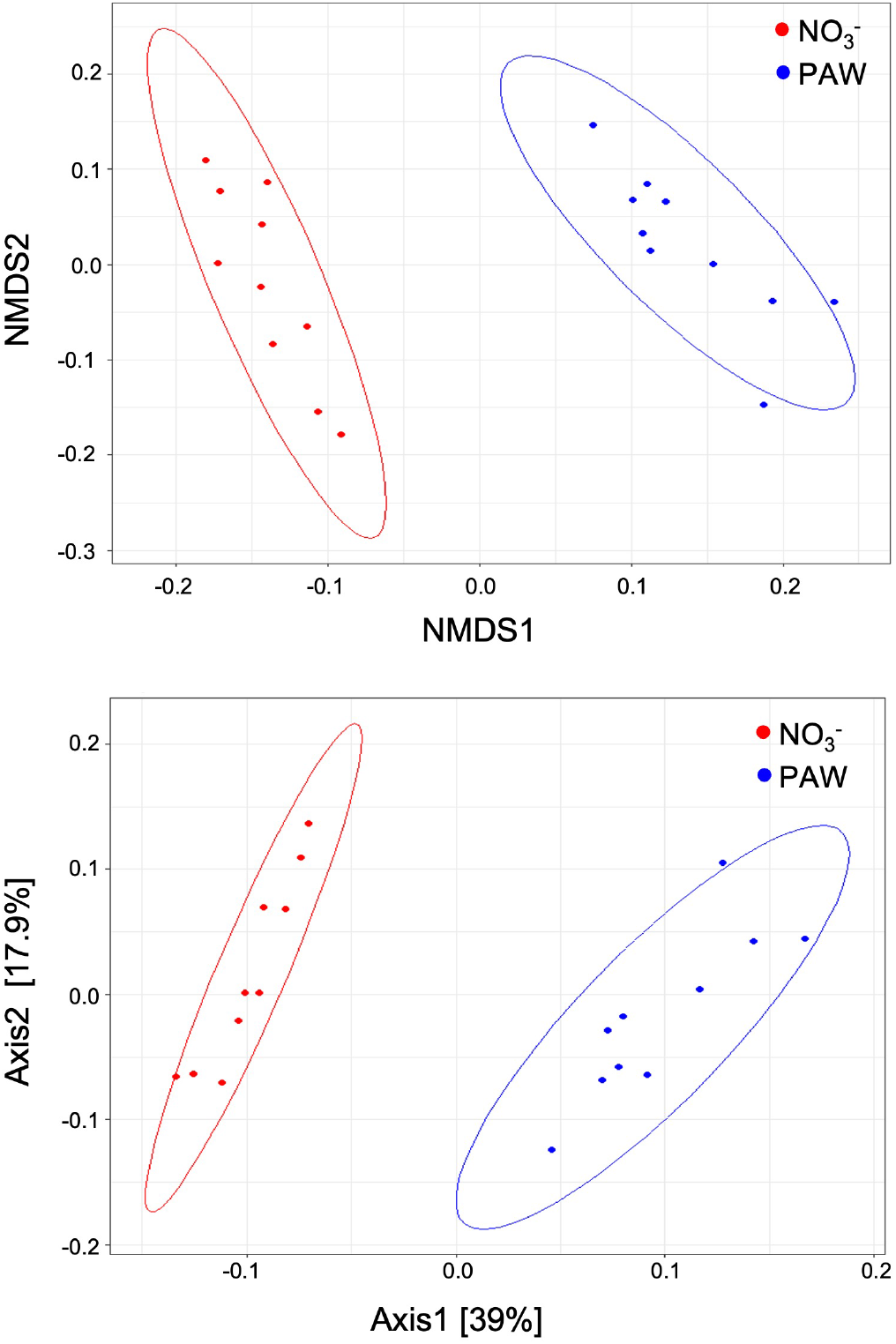
The rhizosphere microbiomes of tomato treated with PAW or the control treatment exhibit dissimilarity in community makeup. Tomato plant rhizosphere microbiomes were identified from plants treated with PAW (6.5 mM NO_3_^-^) or a solution of equivalent NO_3_^-^ concentration for 5 weeks. Amplicon Sequence Variants (ASV) were derived from the sequencing data. Bray-Curtis distances were calculated and plotted via Non-metric Multi-Dimensional Scaling (NMDS) (top) and a Principal Coordinate Analysis (PCoA) (bottom). A residual randomization in permutation procedures (RRPP) analysis was conducted in R to find statistical significance. N= 10 samples per treatment

Taxonomic data was provided from the ASVs processed through the DADA2 pipeline. The Phyloseq R package (McMurdie & Holmes, 2013) was utilized to determine relative abundance of each taxa across treatments. The top 20 taxa were plotted and presented for each sample (Fig. 6). Both PAW and NO_3_^-^ treatments resulted in a similar representation of different bacterial genera. Notably, tomato seedlings treated with PAW had the *Puia* genus represented in the top 20 ASVs while this was observed in only a single seedling treated with the NO_3_^-^ control. While most genera represented amongst the most dominant bacteria are conserved across both treatment groups, the addition of *Puia* coupled with the significant differences in beta diversity calculations suggest that tomato seedlings treated with PAW possess identifiable differences in terms of rhizosphere microbial makeup.

**Figure 6.**
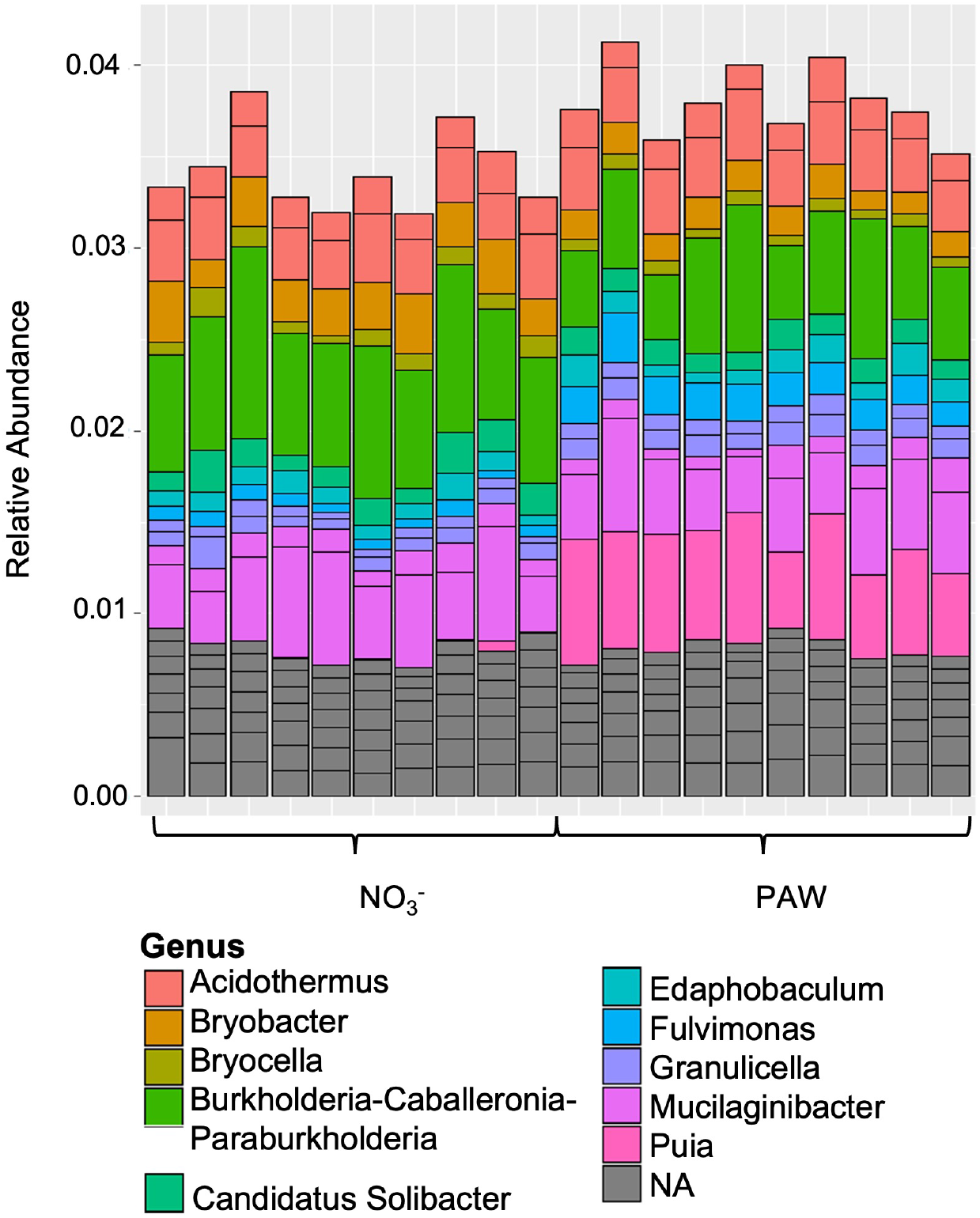
The rhizosphere microbiomes of tomato treated with PAW or the control treatment exhibit similar bacterial genera represented in the 20 most abundant taxa. Tomato plant rhizosphere microbiomes were identified from plants treated with PAW (6.5 mM NO_3_^-^) or a solution of equivalent NO_3_^-^ concentration for 5 weeks and Amplicon Sequence Variants (ASV) were derived from the sequencing data. Taxonomic identities were assigned to ASVs with the Phyloseq R package. The 20 ASVs with the greatest relative abundance from each sample were plotted and grouped by genus.

## DISCUSSION

The root microbiome can be crucial to plant health due to its ability to facilitate nutrient availability and respond to changes in soil conditions (Berendsen et al., 2012; Berg et al., 2014; Hein et al., 2008; Jafariyan & Zarea, 2016; Ney et al., 2019; Richardson & Simpson, 2011; Schneijderberg et al., 2020). The diversity and health of the plant-associated microbiome are contingent on environmental factors (Ai et al., 2015; Berendsen et al., 2012; Berg et al., 2014). Additionally, variations in root exudates promote the association of certain microbes versus others (Badri et al., 2013; Broeckling et al., 2008; Chaparro et al., 2013; Chaparro et al., 2012; Gregory, 2007; Micallef et al., 2009; Sasse et al., 2018; Zhao et al., 2021). In a crop system, the addition of fertilizers impact the state of the plant-associated microbiome (Igiehon & Babalola, 2018; Shi et al., 2020; Zhu et al., 2016) with N fertilizer application rates positively correlated with root exudation rates (Zhu et al., 2016). The effect of fertilizers on microbiome health is relevant when evaluating the efficacy of PAW as an N fertilizer.

PAW is a potential alternative to other less sustainable N-fertilizers. Several studies have shown that PAW has potential enhancements to plant growth when compared to a nitrate-depleted control (Cortese et al., 2021; Cui et al., 2022; Cui et al., 2019; Kučerová et al., 2021; Kučerová et al., 2019; Ndiffo Yemeli et al., 2021; Panngom et al., 2018; Škarpa et al., 2020). From this study, tomato plants supplemented weekly with PAW exhibited comparable growth to those grown with the equivalent NO_3_^-^ control, which indicates that PAW-treatment can substitute a synthetic fertilizer regime. Studies have already captured the potential effects of PAW on plant growth, however PAW’s impacts on the plant-associated microbiome are relatively unexplored. Direct NTP treatment on Arabidopsis seedlings resulted in changes in microbial community make-up with *Mycobacteriaceae* being greatly reduced and *Bacillaceae* abundance being enhanced (Tamošiūnė et al., 2020). Bacteria genera such as *Pseudomonas* and *Paenibacillus* exhibited increased abundance in mature Arabidopsis that grew from NTP-treated seeds (Tamošiūnė et al., 2020), which are known to be beneficial and promote growth (Grady et al., 2016; Santoyo et al., 2012). Changes in microbial community make-up were also observed in seeds directly treated with NTP (Ji et al., 2019). In this study, long-term treatment with PAW and the NO_3_^-^ control did not indicate changes in the functional community makeup of the rhizosphere. Interestingly, bacteria associated with NO_3_^-^ transformations such as those of the Xanthobacteraceae family were represented in the most abundant taxa for both Arabidopsis and tomato which is expected in an N-rich soil environment (Ward, 2013; Yuan et al., 2015).

Establishing potential impacts of PAW treatment on rhizosphere composition and function was the focus of this study in light of extensive research on the rhizosphere microbiome and its important role in maintaining plant health (Hein et al., 2008; Jafariyan & Zarea, 2016; Schneijderberg et al., 2020). In this study, both Arabidopsis and tomato showed similarities in terms of species richness and community make-up between PAW and control-treated plants. This is in contrast to studies in Brassicaceae where changes in N fertilizer source resulted in altered rhizosphere communities by affecting microbe recruitment (Li et al., 2023; O’Brien et al., 2018; Windisch et al., 2021). Additionally, changes in N fertilizers also affected the facilitation of nutrient uptake, including phosphorus via plant-associated microbes in maize (Mang et al., 2023). Even in *S. lycopersicum*, treatments with different synthetic N fertilizers with different chemical composition resulted in an overall enrichment in rhizosphere biodiversity including beneficial *Actinobacteria* (Caradonia et al., 2019). The limited changes in the rhizosphere composition observed in the PAW-treated plants supports use of PAW as an alternative fertilizer, at least for PAWs with minimal ROS.

Depending on varying factors in the plasma-treatment method, ROS are commonly produced in PAW (Cui et al., 2022; Iseni et al., 2016; Kučerová et al., 2019; Ran, Zhou, Wang, et al., 2024). ROS in PAW, such as H_2_O_2_ (Ghimire et al., 2018) or OH radicals (Lamichhane et al., 2019), have shown antimicrobial effects (Graves, 2012; Zhang et al., 2018) leading to PAW being utilized in industrial sterilization (Zhang et al., 2012; Ziuzina et al., 2013) and plasma medicine (Hayashi et al., 2006). Plasma capillary discharge yielding high ROS levels can reduce up to 99.6% *E. coli* load in water (Hong et al., 2010). PAW also has antimicrobial activity against plant pathogens (Ghimire et al., 2024; Ran, Zhou, Dong, et al., 2024). However, PAW without measurable ROS was utilized in this study since ROS in PAW can adversely affect plant growth (Cui et al., 2022; Cui et al., 2019; Ka et al., 2021; Kizer et al., 2025; Zhou et al., 2018). Therefore, other PAW solutions containing ROS may result in changes in the microbiome not captured in this study. Future work should carefully consider the ROS content of PAW when studying plant-associated microbial communities.

This study found that under both treatments, *Burkholderiaceae* bacteria were amongst the most abundant taxa for both Arabidopsis and tomato, specifically those belonging to the combined BCP genus. The BCP group consists of gram-negative bacteria that can either be free-living or endophytic. Bacteria of the *Caballeronia* and *Paraburkhoderia* genera exhibit nitrogen-fixing functions in association with various plant species (Paulitsch et al., 2019; Puri et al., 2020; Suárez-Moreno et al., 2012). In particular, BCP bacteria promote plant growth when associated with Arabidopsis (Adhikari et al., 2019; Huang et al., 2017; Ruiz et al., 2025; K. Wang et al., 2021). NO_3_^-^ replete conditions typically result in higher abundance of this family (Konishi et al., 2017). The *Burkholderiaceae* family is ubiquitous and functionally diverse, with some members isolated as plant pathogens (Adhikari et al., 2019; Jeong et al., 2003; Mannaa et al., 2019), while others act as plant growth promoters (Konishi et al., 2017; O’Sullivan & Mahenthiralingam, 2005; Suárez-Moreno et al., 2012). Given that samples were sequenced at the genera level, an accurate account of the specific species present and their functions is not available in this study. Further studies are needed to determine whether PAW treatment can result in recruitment of specific *Burkholderiaceae* species.

Tomato seedlings treated with PAW exhibited limited differences in rhizosphere microbial composition compared to nitrate treated samples. Specifically, the *Puia* genus was amongst the most abundant ASVs for PAW-treated samples. *Puia* is a relatively unstudied genus of the larger *Chitinophagaceae* family, which was represented by two other genera in both treatments (Supplemental Fig. 3A). Only *Puia dinghuensis* isolated from a forest soil has been characterized (Lv et al., 2017). Other members of the *Chitinophagaceae* have been shown to have beneficial properties for plant growth. For example, these bacteria often produce of plant growth-promoting compounds such as indole-3-acetic acid (Wang et al., 2025), they can break down chitin to release plant-available forms of nutrients (Anzuay et al., 2021; Hui et al., 2020; Jia et al., 2024), and can help control pathogens (Lopez-Nuñez et al., 2025; Nishisaka et al., 2025). It is unclear whether the small changes in *Puia* recruitment would be beneficial to PAW-treated plants.

Overall, this study determined that plants treated with PAW have similar rhizosphere species richness compared to those treated with an inorganic N fertilizer. While limited differences were detected in the respective microbial communities, we found strong similarities when comparing the most abundant taxa. This study constitutes early research on the effects of PAW on the plant-associated microbiome and provides findings that lend support for the use of PAW as a N fertilizer since disruption of plant rhizosphere was not observed for either Arabidopsis or tomato. Further investigation with agricultural soil samples and varying compositions of PAW may provide greater insight as to practical effects of PAW on the plant-associated microbiome.

## MATERIALS AND METHODS

### PAW production

PAW was produced as previously reported (Kizer et al., 2025). using an atmospheric radio frequency (RF) glow discharge plasma. In brief, bulk diH_2_O volumes were exposed to the plasma that was generated by an AE OVAtion 35162 RF at 250 W. Air was flowed down the coaxial electrode, toward the water surface at a rate of ≤ 1 slm. To optimize NO_3_^-^ content, a large external volume of water (≥ 2 L) was circulated through the plasma chamber, which was kept open to improve ventilation, and the distance between the water surface and electrode was set to minimize reflected power (20–40 W reflected at 1.5 cm). Under these conditions, the plasma was consistently able to produce aqueous NO_3_^-^ at a rate of 2 mg/min, and treatment times were adjusted accordingly to achieve the desired concentration for the target volume. The chemical composition was measured colorimetrically and then combined with untreated diH_2_O to obtain the final desired PAW volumes and chemistries. The final PAW would also be tested colorimetrically, to confirm their composition. For each step NO_3_^-^, NO_2_^-^, H_2_O_2_, and NH_4_^+^ were tested for in triplicate using the commercially available Supelco test kits: 1.09713, 1.14776, 1.18789 and 1.14752, respectively. Absorbance values were obtained using an UV-VIS-NIR light source (Ocean Optics DH-2000-BAL) in conjunction with a spectrometer (Ocean Optics QE65 Pro) and a cuvette holder (Ocean Optics CUV-UV). These absorbances were converted to concentrations using stock solution based standard curves prepared in advance. PAW was neutralized with 1M KOH solution to increase the pH to 5.7, a plant-viable pH. Neutralization maintained stability of the solution for storage. PAW was stored in the dark at room temperature for up to 2 weeks, which is unlikely to result in significant changes in nitrate concentration (Risa Vaka et al., 2019; Zhang et al., 2024).

### Plant Material and Growth Conditions

Arabidopsis Columbia 0 (Col-0) ecotype were grown in substrate for this study. Seeds were surface sterilized with 95% ethanol followed by a solution containing 20% commercial bleach and 0.1% Tween 20 (VWR, MFCD00165986). Seeds were rinsed 2-3 times with sterile diH_2_O and then stored at 4°C for 4 days in the dark. Seeds were then plated onto Arabidopsis Growth Media (AGM) containing 0.5 x MS with MES (Murashige & Skoog, 1962, RPI, M70300), 1% sucrose and 4g/L Gelrite (RPI, G35020). Plates were incubated vertically in a growth chamber with 120 µmol/m^2^/s PPFD at 22°C with a 16 h/8 h day/night cycle to promote germination. After transfer to potting substrate, plants were grown on controlled growing benches with LED grow lights under similar controlled conditions as plated plants. PPFD was130 µmol/m^2^/s at plant height. Plants were rotated to new positions regularly on grow benches to reduce the impact of uncontrolled environmental effects. Control solutions were prepared in diH_2_O as 4.8 mM potassium nitrate (KNO_3_^-^) (Caisson Labs, P012) which matched the concentrations of NO_3_^-^ of the PAW.

PAW treatments in substrate were achieved by irrigation of nutrient solutions as follows. A substrate mixture devoid of N was made to avoid any unspecified fertilizer normally present in commercial soil mixes. The substrate mixture contained 45 % peat moss (Premier Peat Moss), 35 % vermiculite (Sta-Green Vermiculite), and 20 % perlite (Aero Soil Perlite) (all measured by volume). Pulverized limestone (Gardenlime Pulverized Dolomitic Limestone) was added to adjust pH to ∼6.0. The substrate was moistened with deionized H_2_O and distributed into 2-inch insert pots (T.O. Plastics, 2401 Standard). Three-day old seedlings previously germinated on sterile AGM media were transferred to each pot of moistened substrate. Pots were incubated at ∼22°C under LED grow lights yielding ∼130 µmol m^-2^ s^-1^ in a 16-hour day cycle. Seedlings were thinned to 1 per pot 3 days after transfer. Seedlings were watered with 50ml per pot diH_2_O twice weekly to keep the plants hydrated, and 50ml per pot of 0.25x Hoagland Solution without N (Bio-World, 30630038) was given to all seedlings biweekly by top irrigation. Specific treatments were provided by top irrigation with 50 ml per pot of either PAW or an equivalent NO_3_^-^ control solution once per week. After 5 weeks of growth, adult Arabidopsis plants were then harvested from the substrate.

Tomato seeds (Burpee, “Big Daddy Hybrid” 69255A) were directly sown to the substrate mix as described above. Two seeds were sown per pot, and excess seedlings were thinned 1 week after sowing. Seedlings were grown for 5 weeks in a growth chamber at 24°C for a 17-hour day and at 16°C for the night under fluorescent grow lights (GE 46761 54W, 417 umol m^-2^ s^-1^). Pots were watered every two days with diH_2_O with increasing volume as needed to maintain consistent substrate moisture. Biweekly, 0.25x Hoagland Solution without N (Bio-World, 30630038) was applied to all pots. Seedlings were treated with either PAW (6.5mM; 400 mg/L NO_3_^-^) or NO_3_^-^ control solution (6.5mM; 400 mg/L NO_3_^-^) starting from week 1. Seedlings were harvested after 5 weeks of growth.

### Rhizosphere Microbiome Analysis

Ten plants were chosen randomly from each treatment, PAW or NO_3_^-^ control, at the end of 5 weeks. Roots were removed from bulk substrate and lightly shaken to remove substrate that was not closely adhered to the root. Then, portions of the remaining substrate were carefully removed with sterile forceps and weighed for harvest. Substrate samples were taken from each individual plant and were not pooled. Bacterial DNA was isolated from approximately 500 mg of substrate sample per plant using the NucleoSpin Soil kit (Takara Bio, 740780). Bacterial 16S rDNA amplicon sequencing was obtained with an Illumina MiSeq (Mr. DNA Lab) using the 515f (Parada) and 806r (Apprill) primers (Apprill et al., 2015; Parada et al., 2016; Walters et al., 2015). Approximately 50,000 reads were per sample were obtained.

Data analysis and processing was all conducted using the R programming language (R-Project). FASTQ files were processed with the DADA2 pipeline (Callahan et al., 2016). Each sample was filtered and trimmed to improve the overall quality of reads. Error-learning, paired-end merging, and chimera removal steps were conducted to maintain quality and accuracy in resulting amplicon sequence variants (ASV) (Callahan et al., 2016). ASVs were further analyzed utilizing the taxonomic tools of the Phyloseq R package (McMurdie & Holmes, 2013) to assign taxonomical IDs to each ASV and calculate both Shannon and Simpson statistics for alpha diversity calculations (Hein et al., 2008; Shannon, 1948). Kruskal-Wallis tests were performed to make pairwise comparisons of both Shannon and Simpson statistics. We visualized all data utilizing the ggplot2 package in R. The Bray-Curtis dissimilarity statistics (Bray & Curtis, 1957) were calculated utilizing the developed ASVs. The resulting Bray-Curtis distance matrix was plotted utilizing the Non-metric MultiDimensional Scaling (NMDS) technique for simple visualization as well as a Principal Coordinate Analysis (PCoA), both utilizing ggplot2 package for R. An RRPP analysis of the distance matrix was used for quantitative confirmation of results.

Lastly, the taxonomic data attributed to the ASVs was processed through the Phyloseq package (McMurdie & Holmes, 2013), and the top 50 most abundant taxa at the genus level in each given sample were found. These taxa were plotted using ggplot2 to visualize the results.

## REFERENCES

Adhikari, B., Adhikari, M., Ghimire, B., Park, G., & Choi, E. H. (2019). Cold Atmospheric Plasma-Activated Water Irrigation Induces Defense Hormone and Gene expression in Tomato seedlings. Scientific Reports, 9(1), 16080. 10.1038/s41598-019-52646-z

Ai, C., Liang, G., Sun, J., Wang, X., He, P., Zhou, W., & He, X. (2015). Reduced dependence of rhizosphere microbiome on plant-derived carbon in 32-year long-term inorganic and organic fertilized soils. Soil Biology and Biochemistry, 80, 70–78. 10.1016/j.soilbio.2014.09.028

Anzuay, M. S., Viso, N. P., Ludueña, L. M., Morla, F. D., Angelini, J. G., & Taurian, T. (2021). Plant beneficial rhizobacteria community structure changes through developmental stages of peanut and maize. Rhizosphere, 19, 100407.

Apprill, A., McNally, S., Parsons, R., & Weber, L. (2015). Minor revision to V4 region SSU rRNA 806R gene primer greatly increases detection of SAR11 bacterioplankton. Aquatic microbial ecology, 75(2), 129–137.

Badri, D. V., Chaparro, J. M., Zhang, R., Shen, Q., & Vivanco, J. M. (2013). Application of natural blends of phytochemicals derived from the root exudates of Arabidopsis to the soil reveal that phenolic-related compounds predominantly modulate the soil microbiome. Journal of Biological Chemistry, 288(7), 4502–4512.

Berendsen, R. L., Pieterse, C. M. J., & Bakker, P. A. H. M. (2012). The rhizosphere microbiome and plant health. Trends in Plant Science, 17(8), 478–486. 10.1016/j.tplants.2012.04.001

Berg, G., Grube, M., Schloter, M., & Smalla, K. (2014). Unraveling the plant microbiome: looking back and future perspectives. Front Microbiol, 5, 148. 10.3389/fmicb.2014.00148

Berrios, L., & Rentsch, J. D. (2022). Linking Reactive Oxygen Species (ROS) to Abiotic and Biotic Feedbacks in Plant Microbiomes: The Dose Makes the Poison. International Journal of Molecular Sciences, 23(8).

Boer, M. D., Santos Teixeira, J., & Ten Tusscher, K. H. (2020). Modeling of Root Nitrate Responses Suggests Preferential Foraging Arises From the Integration of Demand, Supply and Local Presence Signals [Original Research]. Frontiers in Plant Science, 11. 10.3389/fpls.2020.00708

Bray, J. R., & Curtis, J. T. (1957). An ordination of the upland forest communities of southern Wisconsin. Ecological monographs, 27(4), 326–349.

Breidenbach, B., Pump, J., & Dumont, M. G. (2016). Microbial community structure in the rhizosphere of rice plants. Frontiers in microbiology, 6, 1537.

Broeckling, C. D., Broz, A. K., Bergelson, J., Manter, D. K., & Vivanco, J. M. (2008). Root exudates regulate soil fungal community composition and diversity. Applied and environmental microbiology, 74(3), 738–744.

Byrns, B., Wooten, D., Lindsay, A., & Shannon, S. (2012). A VHF driven coaxial atmospheric air plasma: electrical and optical characterization. Journal of Physics D: Applied Physics, 45(19), 195204. 10.1088/0022-3727/45/19/195204

Callahan, B. J., McMurdie, P. J., Rosen, M. J., Han, A. W., Johnson, A. J., & Holmes, S. P. (2016). DADA2: High-resolution sample inference from Illumina amplicon data. Nat Methods, 13(7), 581–583. 10.1038/nmeth.3869

Caradonia, F., Ronga, D., Catellani, M., Giaretta Azevedo, C. V., Terrazas, R. A., Robertson-Albertyn, S., Francia, E., & Bulgarelli, D. (2019). Nitrogen Fertilizers Shape the Composition and Predicted Functions of the Microbiota of Field-Grown Tomato Plants. Phytobiomes Journal, 3(4), 315–325. 10.1094/PBIOMES-06-19-0028-R

Chaparro, J. M., Badri, D. V., & Vivanco, J. M. (2013). Rhizosphere microbiome assemblage is affected by plant development. The ISME Journal, 8(4), 790–803. 10.1038/ismej.2013.196

Chaparro, J. M., Sheflin, A. M., Manter, D. K., & Vivanco, J. M. (2012). Manipulating the soil microbiome to increase soil health and plant fertility. Biology and Fertility of Soils, 48, 489–499.

Chen, J. G., Crooks, R. M., Seefeldt, L. C., Bren, K. L., Bullock, R. M., Darensbourg, M. Y., Holland, P. L., Hoffman, B., Janik, M. J., Jones, A. K., Kanatzidis, M. G., King, P., Lancaster, K. M., Lymar, S. V., Pfromm, P., Schneider, W. F., & Schrock, R. R. (2018). Beyond fossil fuel–driven nitrogen transformations. Science, 360(6391), eaar6611. 10.1126/science.aar6611

Claesson, M. J., Wang, Q., O’Sullivan, O., Greene-Diniz, R., Cole, J. R., Ross, R. P., & O’Toole, P. W. (2010). Comparison of two next-generation sequencing technologies for resolving highly complex microbiota composition using tandem variable 16S rRNA gene regions. Nucleic acids research, 38(22), e200–e200.

Cortese, E., Settimi, A. G., Pettenuzzo, S., Cappellin, L., Galenda, A., Famengo, A., Dabalà, M., Antoni, V., & Navazio, L. (2021). Plasma-Activated Water Triggers Rapid and Sustained Cytosolic Ca2+ Elevations in Arabidopsis thaliana. Plants, 10(11).

Cui, D., Yin, Y., Sun, H., Wang, X., Zhuang, J., Wang, L., Ma, R., & Jiao, Z. (2022). Regulation of cellular redox homeostasis in Arabidopsis thaliana seedling by atmospheric pressure cold plasma-generated reactive oxygen/nitrogen species. Ecotoxicology and Environmental Safety, 240, 113703. 10.1016/j.ecoenv.2022.113703

Cui, D., Yin, Y., Wang, J., Wang, Z., Ding, H., Ma, R., & Jiao, Z. (2019). Research on the Physio-Biochemical Mechanism of Non-Thermal Plasma-Regulated Seed Germination and Early Seedling Development in Arabidopsis. Front Plant Sci, 10, 1322. 10.3389/fpls.2019.01322

Ghimire, B., Lee, G. J., Mumtaz, S., & Choi, E. H. (2018). Scavenging effects of ascorbic acid and mannitol on hydroxyl radicals generated inside water by an atmospheric pressure plasma jet. AIP Advances, 8(7). 10.1063/1.5037125

Ghimire, B., Pendyala, B., Patras, A., & Baysal-Gurel, F. (2024). Effect of Plasma-Activated Water (PAW) Generated Using Non-Thermal Atmospheric Plasma on Phytopathogenic Bacteria. Plant Disease. 10.1094/PDIS-05-24-0957-SC

Gilbert, J., Jansson, J., & Knight, R. (2014). The Earth Microbiome project: successes and. Turnbaugh PJ, Walters WA, Widmann J, Yatsunenko T, Zaneveld J, Knight R: QIIME.

Grady, E. N., MacDonald, J., Liu, L., Richman, A., & Yuan, Z.-C. (2016). Current knowledge and perspectives of Paenibacillus: a review. Microbial cell factories, 15, 1–18.

Graves, D. B. (2012). The emerging role of reactive oxygen and nitrogen species in redox biology and some implications for plasma applications to medicine and biology. Journal of Physics D: Applied Physics, 45(26), 263001.

Gregory, P. (2007). Plant roots. Wiley Online Library.

Hayashi, N., Guan, W., Tsutsui, S., Tomari, T., & Hanada, Y. (2006). Sterilization of Medical Equipment Using Radicals Produced by Oxygen/Water Vapor RF Plasma. Japanese Journal of Applied Physics, 45(10S), 8358. 10.1143/JJAP.45.8358

Hein, J. W., Wolfe, G. V., & Blee, K. A. (2008). Comparison of Rhizosphere Bacterial Communities in Arabidopsis thaliana Mutants for Systemic Acquired Resistance. Microbial Ecology, 55(2), 333–343. 10.1007/s00248-007-9279-1

Hill, T. C. J., Walsh, K. A., Harris, J. A., & Moffett, B. F. (2003). Using ecological diversity measures with bacterial communities. FEMS Microbiology Ecology, 43(1), 1–11. 10.1111/j.1574-6941.2003.tb01040.x

Hong, S.-H., Bunge, J., Jeon, S.-O., & Epstein, S. S. (2006). Predicting microbial species richness. Proceedings of the National Academy of Sciences, 103(1), 117–122.

Hong, Y. C., Park, H. J., Lee, B. J., Kang, W.-S., & Uhm, H. S. (2010). Plasma formation using a capillary discharge in water and its application to the sterilization of E. coli. Physics of Plasmas, 17(5). 10.1063/1.3418371

Huang, X.-F., Zhou, D., Lapsansky, E. R., Reardon, K. F., Guo, J., Andales, M. J., Vivanco, J. M., & Manter, D. K. (2017). Mitsuaria sp. and Burkholderia sp. from Arabidopsis rhizosphere enhance drought tolerance in Arabidopsis thaliana and maize (Zea mays L.). Plant and Soil, 419(1), 523–539. 10.1007/s11104-017-3360-4

Hui, C., Jiang, H., Liu, B., Wei, R., Zhang, Y., Zhang, Q., Liang, Y., & Zhao, Y. (2020). Chitin degradation and the temporary response of bacterial chitinolytic communities to chitin amendment in soil under different fertilization regimes. Science of the Total Environment, 705, 136003.

Igiehon, N. O., & Babalola, O. O. (2018). Rhizosphere Microbiome Modulators: Contributions of Nitrogen Fixing Bacteria towards Sustainable Agriculture. International Journal of Environmental Research and Public Health, 15(4).

Iseni, S., Bruggeman, P. J., Weltmann, K.-D., & Reuter, S. (2016). Nitrogen metastable (N2(A3Σu+)) in a cold argon atmospheric pressure plasma jet: Shielding and gas composition. Applied Physics Letters, 108(18), 184101. 10.1063/1.4948535

Jafariyan, T., & Zarea, M. J. (2016). Hydrogen peroxide affects plant growth promoting effects of Azospirillum. Journal of Crop Science and Biotechnology, 19(2), 167–175. 10.1007/s12892-015-0127-4

Jeong, Y., Kim, J., Kim, S., Kang, Y., Nagamatsu, T., & Hwang, I. (2003). Toxoflavin produced by Burkholderia glumae causing rice grain rot is responsible for inducing bacterial wilt in many field crops. Plant Disease, 87(8), 890–895.

Ji, S.-H., Kim, J.-S., Lee, C.-H., Seo, H.-S., Chun, S.-C., Oh, J., Choi, E.-H., & Park, G. (2019). Enhancement of vitality and activity of a plant growth-promoting bacteria (PGPB) by atmospheric pressure non-thermal plasma. Scientific Reports, 9(1), 1044. 10.1038/s41598-018-38026-z

Jia, X., Shang, H., Chen, Y., Lin, M., Wei, Y., Li, Y., Li, R., Dong, P., Chen, Y., & Zhang, Y. (2024). Improved bacterial composition and co-occurrence patterns of rhizosphere increased nutrient uptake and grain yield through cultivars mixtures in maize. Science of the Total Environment, 926, 172102.

Ka, D. H., Priatama, R. A., Park, J. Y., Park, S. J., Kim, S. B., Lee, I. A., & Lee, Y. K. (2021). Plasma-Activated Water Modulates Root Hair Cell Density via Root Developmental Genes in Arabidopsis thaliana L. Applied Sciences, 11(5).

Kizer, J., Robinson, C., Lucas, T. k., Shannon, S., Hernandez, R., Stapelmann, K., & Rojas-Pierce, M. (2025). Non-thermal plasma activated water is an effective nitrogen fertilizer alternative for Arabidopsis thaliana. PLoS ONE, in print.

Konishi, N., Okubo, T., Yamaya, T., Hayakawa, T., & Minamisawa, K. (2017). Nitrate Supply-Dependent Shifts in Communities of Root-Associated Bacteria in <i>Arabidopsis</i>. Microbes and Environments, 32(4), 314–323. 10.1264/jsme2.ME17031

Kučerová, K., Henselová, M., Slováková, Ľ., Bačovčinová, M., & Hensel, K. (2021). Effect of Plasma Activated Water, Hydrogen Peroxide, and Nitrates on Lettuce Growth and Its Physiological Parameters. Applied Sciences, 11(5).

Kučerová, K., Henselová, M., Slováková, Ľ., & Hensel, K. (2019). Effects of plasma activated water on wheat: Germination, growth parameters, photosynthetic pigments, soluble protein content, and antioxidant enzymes activity. Plasma Processes and Polymers, 16(3), 1800131. 10.1002/ppap.201800131

Lamichhane, P., Ghimire, B., Mumtaz, S., Paneru, R., Ki, S. H., & Choi, E. H. (2019). Control of hydrogen peroxide production in plasma activated water by utilizing nitrification. Journal of Physics D: Applied Physics, 52(26), 265206. 10.1088/1361-6463/ab16a9

Lassaletta, L., Billen, G., Grizzetti, B., Anglade, J., & Garnier, J. (2014). 50 year trends in nitrogen use efficiency of world cropping systems: the relationship between yield and nitrogen input to cropland. Environmental Research Letters, 9(10), 105011. 10.1088/1748-9326/9/10/105011

Li, Y., Vail, S. L., Arcand, M. M., & Helgason, B. L. (2023). Contrasting Nitrogen Fertilization and Brassica napus (Canola) Variety Development Impact Recruitment of the Root-Associated Microbiome. Phytobiomes Journal, 7(1), 125–137. 10.1094/PBIOMES-07-22-0045-R

Lindsay, A., Byrns, B., King, W., Andhvarapou, A., Fields, J., Knappe, D., Fonteno, W., & Shannon, S. (2014). Fertilization of radishes, tomatoes, and marigolds using a large-volume atmospheric glow discharge. Plasma Chemistry and Plasma Processing, 34, 1271–1290.

Lopez-Nuñez, R., Prieto-Rubio, J., Bautista, I., Lidon-Cerezuela, A. L., Valverde-Urrea, M., Lopez-Moya, F., & Lopez-Llorca, L. V. (2025). Chitosan reduces naturally occurring plant pathogenic fungi and increases nematophagous fungus Purpureocillium in soil under field conditions. Frontiers in Agronomy, 6, 1502402.

Lv, Y.-y., Gao, Z.-h., Xia, F., Chen, M.-h., & Qiu, L.-h. (2017). Puia dinghuensis gen. nov., sp. nov., isolated from monsoon evergreen broad-leaved forest soil. International Journal of Systematic and Evolutionary Microbiology, 67(11), 4639–4645. 10.1099/ijsem.0.002346

Mang, M., Maywald, N. J., Li, X., Ludewig, U., & Francioli, D. (2023). Nitrogen Fertilizer Type and Genotype as Drivers of P Acquisition and Rhizosphere Microbiota Assembly in Juvenile Maize Plants. Plants, 12(3).

Mannaa, M., Park, I., & Seo, Y.-S. (2019). Genomic Features and Insights into the Taxonomy, Virulence, and Benevolence of Plant-Associated Burkholderia Species. International Journal of Molecular Sciences, 20(1), 121. http://www.mdpi.com/1422-0067/20/1/121

McMurdie, P. J., & Holmes, S. (2013). phyloseq: an R package for reproducible interactive analysis and graphics of microbiome census data. PloS one, 8(4), e61217.

Menegat, S., Ledo, A., & Tirado, R. (2022). Greenhouse gas emissions from global production and use of nitrogen synthetic fertilisers in agriculture. Scientific Reports, 12(1), 1–13.

Micallef, S. A., Shiaris, M. P., & Colón-Carmona, A. (2009). Influence of Arabidopsis thaliana accessions on rhizobacterial communities and natural variation in root exudates. Journal of experimental botany, 60(6), 1729–1742.

Ndijo Yemeli, G. B., Švubová, R., Kostolani, D., Kyzek, S., & Machala, Z. (2021). The effect of water activated by nonthermal air plasma on the growth of farm plants: Case of maize and barley. Plasma Processes and Polymers, 18(1), 2000205. 10.1002/ppap.202000205

Ney, L., Franklin, D., Mahmud, K., Cabrera, M., Hancock, D., Habteselassie, M., Newcomer, Q., & Fatzinger, B. (2019). Rebuilding Soil Ecosystems for Improved Productivity in Biosolarized Soils. International Journal of Agronomy, 2019(1), 5827585. 10.1155/2019/5827585

Nishisaka, C. S., Quevedo, H. D., Ventura, J. P., Andreote, F. D., Mauchline, T. H., & Mendes, R. (2025). Soil Microbial Diversity: A Key Factor in Pathogen Suppression and Inoculant Performance. Available at SSRN 5154685.

O’Brien, F. J. M., Dumont, M. G., Webb, J. S., & Poppy, G. M. (2018). Rhizosphere Bacterial Communities Differ According to Fertilizer Regimes and Cabbage (Brassica oleracea var. capitata L.) Harvest Time, but Not Aphid Herbivory [Original Research]. Frontiers in microbiology, 9. 10.3389/fmicb.2018.01620

O’Sullivan, L. A., & Mahenthiralingam, E. (2005). Biotechnological potential within the genus Burkholderia. Letters in applied microbiology, 41(1), 8–11.

Panngom, K., Chuesaard, T., Tamchan, N., Jiwchan, T., Srikongsritong, K., & Park, G. (2018). Comparative assessment for the effects of reactive species on seed germination, growth and metabolisms of vegetables. Scientia Horticulturae, 227, 85–91. 10.1016/j.scienta.2017.09.026

Parada, A. E., Needham, D. M., & Fuhrman, J. A. (2016). Every base matters: assessing small subunit rRNA primers for marine microbiomes with mock communities, time series and global field samples. Environmental microbiology, 18(5), 1403–1414.

Paulitsch, F., Dall’Agnol, R. F., Delamuta, J. R. M., Ribeiro, R. A., da Silva Batista, J.S., & Hungria, M. (2019). Paraburkholderia guartelaensis sp. nov., a nitrogen-fixing species isolated from nodules of Mimosa gymnas in an ecotone considered as a hotspot of biodiversity in Brazil. Archives of Microbiology, 201(10), 1435–1446. 10.1007/s00203-019-01714-z

Puri, A., Padda, K. P., & Chanway, C. P. (2020). Can naturally-occurring endophytic nitrogen-fixing bacteria of hybrid white spruce sustain boreal forest tree growth on extremely nutrient-poor soils? Soil Biology and Biochemistry, 140, 107642. 10.1016/j.soilbio.2019.107642

Quast, C., Pruesse, E., Yilmaz, P., Gerken, J., Schweer, T., Yarza, P., Peplies, J., & Glöckner, F. O. (2012). The SILVA ribosomal RNA gene database project: improved data processing and web-based tools. Nucleic acids research, 41(D1), D590-D596. 10.1093/nar/gks1219

Ran, C., Zhou, X., Dong, P., Liu, K., & Ostrikov, K. (2024). Ultralong-lasting plasma-activated water inhibits pathogens and improves plant disease resistance in soft rot-infected hydroponic lettuce. Plasma Processes and Polymers, 21(8), e2400039. 10.1002/ppap.202400039

Ran, C., Zhou, X., Wang, Z., Liu, K., & Ostrikov, K. (2024). Ultralong-lasting plasma-activated water: production and control mechanisms. Plasma Sources Science and Technology, 33(1), 015009. 10.1088/1361-6595/ad1b6c

Richardson, A. E., & Simpson, R. J. (2011). Soil microorganisms mediating phosphorus availability update on microbial phosphorus. Plant physiology, 156(3), 989–996.

Risa Vaka, M., Sone, I., García Álvarez, R., Walsh, J. L., Prabhu, L., Sivertsvik, M., & Noriega Fernández, E. A.-O. (2019). Towards the Next-Generation Disinfectant: Composition, Storability and Preservation Potential of Plasma Activated Water on Baby Spinach Leaves. LID -10.3390/foods8120692 [doi] LID - 692. (2304-8158 (Print)).

Ruiz, D., Céspedes-Bernal, N., Vega, A., Ledger, T., González, B., & Poupin, M. J. (2025). Nitrogen-modulated effects of the diazotrophic bacterium Cupriavidus taiwanensis on the non-nodulating plant Arabidopsis thaliana. Plant and Soil, 506(1), 819–837. 10.1007/s11104-024-06736-1

Santoyo, G., Orozco-Mosqueda, M. d. C., & Govindappa, M. (2012). Mechanisms of biocontrol and plant growth-promoting activity in soil bacterial species of Bacillus and Pseudomonas: a review. Biocontrol Science and Technology, 22(8), 855–872.

Sasse, J., Martinoia, E., & Northen, T. (2018). Feed your friends: do plant exudates shape the root microbiome? Trends in Plant Science, 23(1), 25–41.

Schneijderberg, M., Cheng, X., Franken, C., de Hollander, M., van Velzen, R., Schmitz, L., Heinen, R., Geurts, R., van der Putten, W. H., Bezemer, T. M., & Bisseling, T. (2020). Quantitative comparison between the rhizosphere effect of Arabidopsis thaliana and co-occurring plant species with a longer life history. The ISME Journal, 14(10), 2433–2448. 10.1038/s41396-020-0695-2

Shannon, C. E. (1948). A mathematical theory of communication. The Bell system technical journal, 27(3), 379–423.

Shi, Y., Ziadi, N., Hamel, C., Bélanger, G., Abdi, D., Lajeunesse, J., Lafond, J., Lalande, R., & Shang, J. (2020). Soil microbial biomass, activity and community structure as affected by mineral phosphorus fertilization in grasslands. Applied Soil Ecology, 146, 103391.

Škarpa, P., Klofáč, D., Krčma, F., Šimečková, J., & Kozáková, Z. (2020). Effect of Plasma Activated Water Foliar Application on Selected Growth Parameters of Maize (Zea mays L.). Water, 12(12).

Suárez-Moreno, Z. R., Caballero-Mellado, J., Coutinho, B. G., Mendonça-Previato, L., James, E. K., & Venturi, V. (2012). Common features of environmental and potentially beneficial plant-associated Burkholderia. Microbial Ecology, 63, 249–266.

Tamošiūnė, I., Gelvonauskienė, D., Ragauskaitė, L., Koga, K., Shiratani, M., & Baniulis, D. (2020). Cold plasma treatment of Arabidopsis thaliana (L.) seeds modulates plant-associated microbiome composition. Applied Physics Express, 13(7), 076001. 10.35848/1882-0786/ab9712

Truyens, S., Weyens, N., Cuypers, A., & Vangronsveld, J. (2013). Changes in the population of seed bacteria of transgenerationally Cd-exposed A rabidopsis thaliana. Plant Biology, 15(6), 971–981.

Truyens, S., Weyens, N., Cuypers, A., & Vangronsveld, J. (2015). Bacterial seed endophytes: genera, vertical transmission and interaction with plants. Environmental Microbiology Reports, 7(1), 40–50. 10.1111/1758-2229.12181

Tsukagoshi, H., Busch, W., & Benfey, P. N. (2010). Transcriptional Regulation of ROS Controls Transition from Proliferation to Differentiation in the Root. Cell, 143(4), 606–616. 10.1016/j.cell.2010.10.020

Walters, W., Hyde, E. R., Berg-Lyons, D., Ackermann, G., Humphrey, G., Parada, A., Gilbert, J. A., Jansson, J. K., Caporaso, J. G., Fuhrman, J. A., Apprill, A., & Knight, R. (2015). Improved Bacterial 16S rRNA Gene (V4 and V4-5) and Fungal Internal Transcribed Spacer Marker Gene Primers for Microbial Community Surveys. LID - e00009-15. (2379-5077 (Print)).

Wang, K., Wu, Y., Ye, M., Yang, Y., Asiegbu, F. O., Overmyer, K., Liu, S., & Cui, F. (2021). Comparative genomics reveals potential mechanisms of plant beneficial effects of a novel bamboo-endophytic bacterial isolate Paraburkholderia sacchari Suichang626. Frontiers in microbiology, 12, 686998.

Wang, M., Khan, M. A., Mohsin, I., Wicks, J., Ip, A. H., Sumon, K. Z., Dinh, C.-T., Sargent, E. H., Gates, I. D., & Kibria, M. G. (2021). Can sustainable ammonia synthesis pathways compete with fossil-fuel based Haber–Bosch processes? [10.1039/D0EE03808C]. Energy & Environmental Science, 14(5), 2535–2548. 10.1039/D0EE03808C

Wang, Y., Zhang, Y., Song, T., Wang, W., Dong, J., Wei, X., & Lin, Y. (2025). Recruitment of beneficial microorganisms by biogas fertilizer enhances the yield and quality of Gynostemma pentaphyllum. Plant and Soil, 1–16.

Ward, B. B. (2013). Nitrification☆. In B. Fath (Ed.), Encyclopedia of Ecology (Second Edition) (pp. 351–358). Elsevier. 10.1016/B978-0-12-409548-9.00697-7

Windisch, S., Sommermann, L., Babin, D., Chowdhury, S. P., Grosch, R., Moradtalab, N., Walker, F., Höglinger, B., El-Hasan, A., Armbruster, W., Nesme, J., Sørensen, S. J., Schellenberg, I., Geistlinger, J., Smalla, K., Rothballer, M., Ludewig, U., & Neumann, G. (2021). Impact of Long-Term Organic and Mineral Fertilization on Rhizosphere Metabolites, Root–Microbial Interactions and Plant Health of Lettuce [Original Research]. Frontiers in microbiology, 11. 10.3389/fmicb.2020.597745

Yuan, J., Chaparro, J. M., Manter, D. K., Zhang, R., Vivanco, J. M., & Shen, Q. (2015). Roots from distinct plant developmental stages are capable of rapidly selecting their own microbiome without the influence of environmental and soil edaphic factors. Soil Biology and Biochemistry, 89, 206–209. 10.1016/j.soilbio.2015.07.009

Zhang, H., Momna, R., Moutong, C., Jialong, G., Qinxiu, S., Qiuyu, X., Zefu, W., Zongyuan, H., Shucheng, L., & and Wei, S. (2024). Study on the detection of active components in plasma-activated water and its storage stability. CyTA - Journal of Food, 22(1), 2386417. 10.1080/19476337.2024.2386417

Zhang, X., Liu, D., Wang, H., Liu, L., Wang, S., & Yang, S.-z. (2012). Highly Effective Inactivation of Pseudomonas sp HB1 in Water By Atmospheric Pressure Microplasma Jet Array. Plasma Chemistry and Plasma Processing, 32(5), 949–957. 10.1007/s11090-012-9389-5

Zhang, X., Zhou, R., Bazaka, K., Liu, Y., Zhou, R., Chen, G., Chen, Z., Liu, Q., Yang, S., & Ostrikov, K. (2018). Quantification of plasma produced OH radical density for water sterilization. Plasma Processes and Polymers, 15(6), 1700241. 10.1002/ppap.201700241

Zhang, Y., Xia, C., Zhang, X., Sha, Y., Feng, G., & Gao, Q. (2022). Quantifying the relationships of soil properties and crop growth with yield in a NPK fertilizer application maize field. Computers and Electronics in Agriculture, 198, 107011. 10.1016/j.compag.2022.107011

Zhao, M., Zhao, J., Yuan, J., Hale, L., Wen, T., Huang, Q., Vivanco, J. M., Zhou, J., Kowalchuk, G. A., & Shen, Q. (2021). Root exudates drive soil-microbe-nutrient feedbacks in response to plant growth. Plant, Cell & Environment, 44(2), 613–628. 10.1111/pce.13928

Zhou, L., Hou, H., Yang, T., Lian, Y., Sun, Y., Bian, Z., & Wang, C. (2018). Exogenous hydrogen peroxide inhibits primary root gravitropism by regulating auxin distribution during Arabidopsis seed germination. Plant Physiology and Biochemistry, 128, 126–133. 10.1016/j.plaphy.2018.05.014

Zhu, S., Vivanco, J. M., & Manter, D. K. (2016). Nitrogen fertilizer rate affects root exudation, the rhizosphere microbiome and nitrogen-use-efficiency of maize. Applied Soil Ecology, 107, 324–333. 10.1016/j.apsoil.2016.07.009

Ziuzina, D., Patil, S., Cullen, P. J., Keener, K. M., & Bourke, P. (2013). Atmospheric cold plasma inactivation of Escherichia coli in liquid media inside a sealed package. Journal of Applied Microbiology, 114(3), 778–787. 10.1111/jam.12087

