## Supplemental Material for "The Effect of Plasma Activated Water on the Rhizosphere Composition of Arabidopsis and *Solanum lycopersicum*"

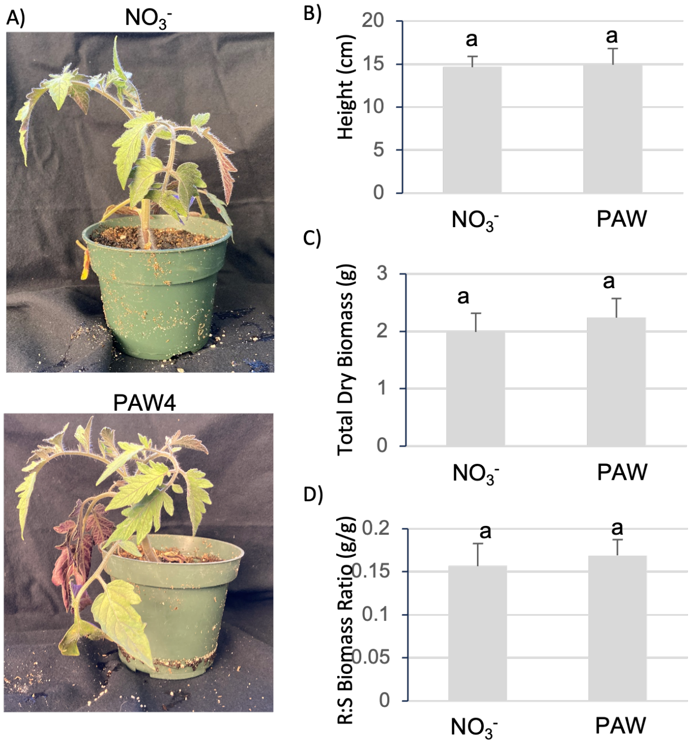


**Supplemental Figure 1. PAW-treated tomato show comparable growth to plants fertilized by the $NO_3^-$ control**

**A)** Tomato plants were grown in substrate and watered weekly with PAW (6.5 mM $NO_3^-$) or a solution of equivalent $NO_3^-$ concentration for 5 weeks. **B-D)** Plants were harvested after 5 weeks of treatment, and shoots and roots were used to measure plant height (B), total dry biomass (C), and root:shoot ratio D). N = 20 plants per treatment. Significance denotes $p < 0.05$ in Student's T-test.

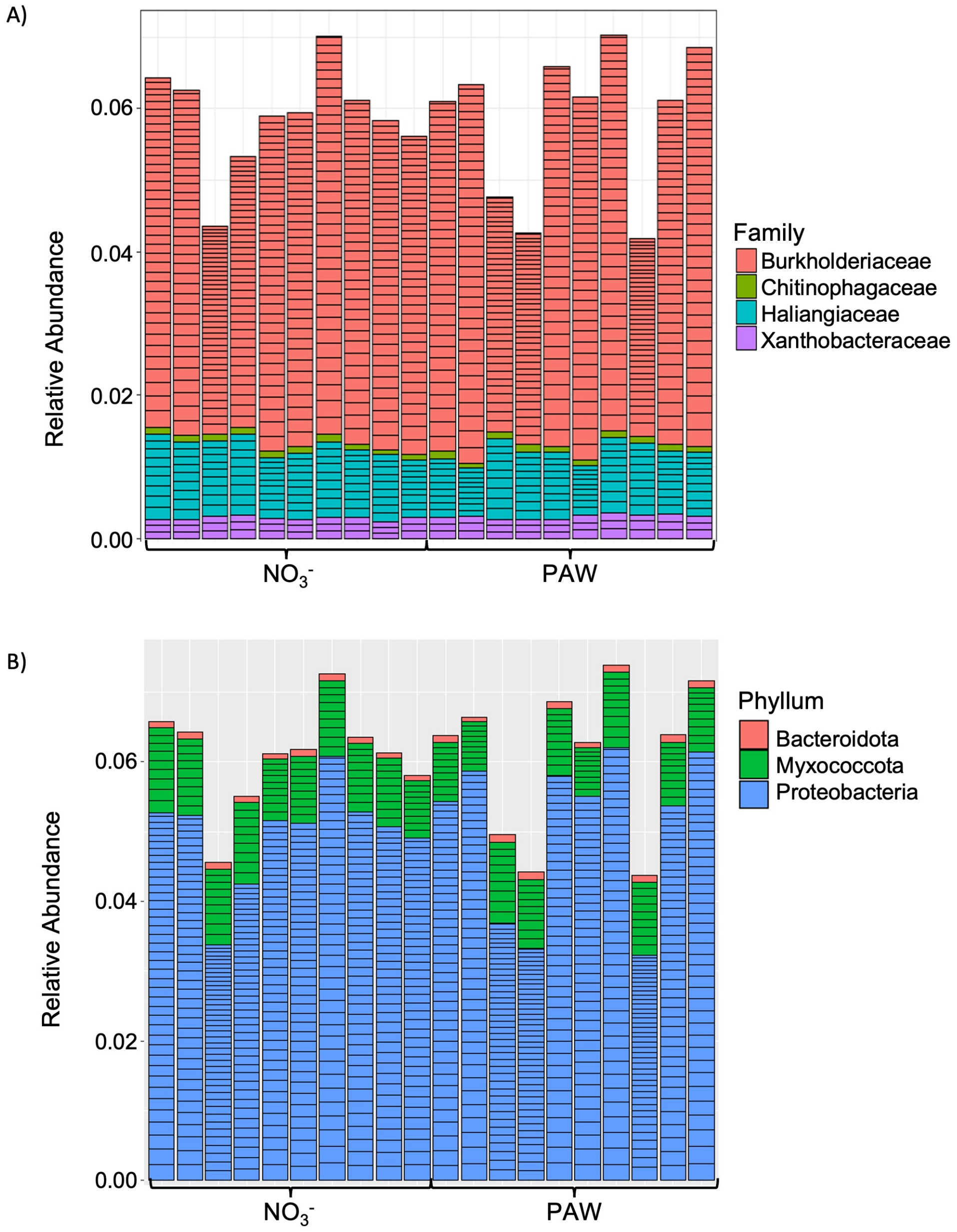


**Supplemental Figure 2. The rhizosphere microbiomes of Arabidopsis treated with PAW or the control treatment exhibit similar bacterial genera and phyla represented in the 50 most abundant taxa.**

Arabidopsis rhizosphere microbiomes were identified from plants treated with PAW (4.8 mM $NO_3^-$) or a solution of equivalent $NO_3^-$ concentration for 5 weeks and Amplicon Sequence Variants (ASV) were derived from the sequencing data. Taxonomic identities were assigned to ASVs with the Phyloseq R package. The 50 ASVs with the greatest relative abundance from each sample were plotted and grouped by **A)** genera and **B**) phyla.

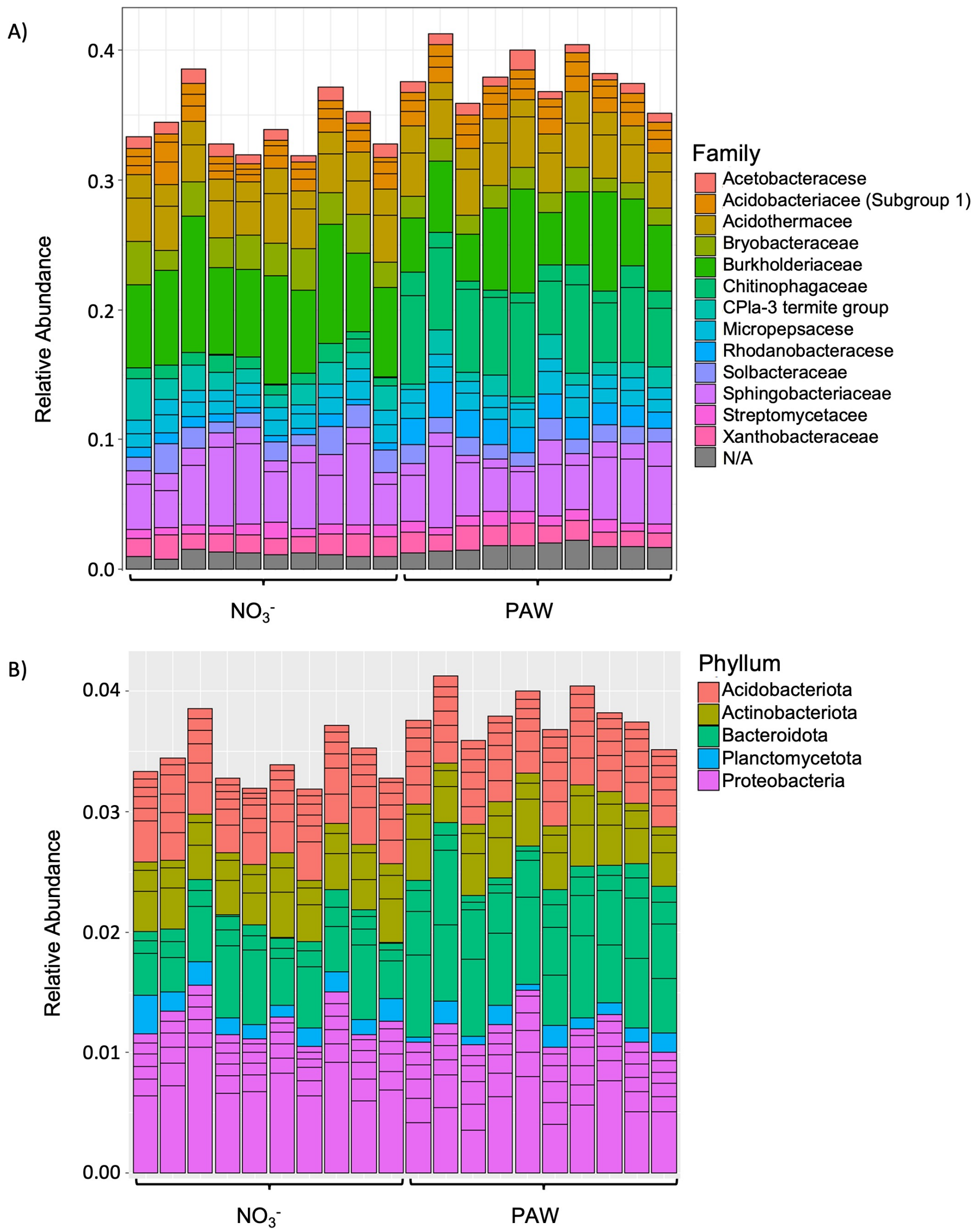


**Supplemental Figure 3. The rhizosphere microbiomes of tomato treated with PAW or the control treatment exhibit similar bacterial families represented in the 20 most abundant taxa.**

Tomato plant rhizosphere microbiomes were identified from plants treated with PAW (6.5 mM $NO_3^-$) or a solution of equivalent $NO_3^-$ concentration for 5 weeks and Amplicon Sequence Variants (ASV) were derived from the sequencing data. Taxonomic identities were assigned to ASVs with the Phyloseq R package. The 20 ASVs with the greatest relative abundance from each sample were plotted and grouped by **A)** genera and **B**) phyla.
